# Combining free energy simulations and NMR chemical-shift perturbation to identify transient cation-*π* contacts in proteins

**DOI:** 10.1101/793984

**Authors:** André A. O. Reis, Raphael S. R. Sayegh, Sandro R. Marana, Guilherme M. Arantes

## Abstract

Flexible protein regions containing cationic and aromatic side-chains exposed to solvent may form transient cation-*π* interactions with structural and functional roles. To evaluate their stability and identify important intramolecular cation-*π* contacts, a combination of free energy profiles estimated from umbrella sampling with molecular dynamics simulations and chemical shift perturbations (CSP) obtained from NMR experiments is applied here to the complete catalytic domain of human phosphatase Cdc25B. This protein is a good model system for transient cation-*π* interactions as it contains only one Trp residue (W550) in the disordered C-terminal segment and a total of 17 Arg residues, many exposed to solvent. Eight putative Arg-Trp pairs were simulated here. Only R482 and R544 show bound profiles corresponding to important transient cation-*π* interactions, while the others have dissociative or almost flat profiles. These results are corroborated by CSP analysis of three Cdc25B point mutants (W550A, R482A and R544A) disrupting cation-*π* contacts. The proposed validation of statistically representative molecular simulations by NMR spectroscopy could be applied to identify transient contacts of proteins in general but carefully, as NMR chemical shifts are sensitive to changes in both molecular contacts and conformational distributions.

## 1 Introduction

Proteins are stabilized by multiple noncovalent intramolecular contacts such as hydrogen bonds, hydrophobic contacts and ion pairs. Cation-*π* interactions formed between cationic residues (in Arg and Lys) and aromatic rings (in Phe, Tyr and Trp) have comparable interaction energies mainly of electrostatic origin^1^ and are easy to recognize when buried in folded proteins.^2^ Cation-*π* contacts may also be observed between side-chains exposed to solvent due to the competition of aromatic rings with water in binding cations in aqueous environments.^1,3,4^

The lifetime of a non-covalent contact in aqueous solution is determined by its stability or free energy, and weak contacts will survive briefly due to structural fluctuations and water competition.^5^ Flexible protein regions such as side-chains exposed to solvent, loops and intrinsically disordered segments should have their structure represented by a conformational ensemble in which the stability of a possible non-covalent contact is proportional to its population or statistical importance in the configurational distribution.^6^ However, ensembles commonly obtained from trajectories of molecular dynamics (MD) simulation often sample the underlying equilibrium distribution inappropriately or incompletely.^7^ Not seldom, a single observation of a transient contact during a simulation is taken as evidence of its structural or functional role^8–10^ but without statistical significance.

Here, the statistical importance of cation-*π* interactions formed between protein side-chains is investigated with a combination of molecular simulation and experimental spectroscopy. Enhanced sampling free energy simulations are employed. In principle, these may lead to correct contact distributions if a meaningful collective or reaction coordinate describing the transient interaction is chosen and orthogonal degrees of freedom are properly sampled.^7^ But, given the approximate description of interaction energies used in biomolecular simulations and finite sampling available in practice, an experimental validation of putative transient contacts will help to confirm the computational predictions.

Nuclear magnetic resonance (NMR) is exquisitely sensitive to the nuclei chemical environment and is routinely used to determine intramolecular contacts of proteins in solution.^11^ For instance, cation-*π* interactions will shift Arg and Lys side-chain resonances upfield because of diamagnetic anisotropic shielding from a nearby indole ring in Trp.^4^ But NMR chemical shifts also report on the distribution of protein conformations. The nuclear Overhauser effect (NOE) is frequently used to discriminate shifts induced by molecular contact from those of conformational origin.^11^ Here, only standard ^1^H-^15^N HSQC spectra were explored to avoid additional NMR experiments or extensive assignments of side-chain resonances,^12^ so that the proposed experimental validation is kept as simple and general as possible. It should be noted that other spectroscopical methods such as visible absorbance and fluorescence have recently been used to identify cation-*π* interactions in solution^13^ and could also be employed to validate simulations of transient contacts if enough spectral resolution is available.

The complete catalytic domain of human phosphatase Cdc25B was studied as a model system.^14,15^ This enzyme catalyzes the de-phosphorylation of pTyr and pThr from other peptides^16–19^ and has already been investigated by solution NMR.^20,21^ It contains only one Trp residue (W550) located in the disordered C-terminus,^21,22^ as well as a total of 17 Arg (5 in the disordered C-terminus) and 13 Lys. Many of these Arg side-chains are exposed on the protein surface near the P-loop catalytic site and W550 (Fig. 1). The large number of cationic residues in Cdc25B may be related to anionic substrate recognition and product release.^15^ Yet, multiple Trp-Arg contacts were previously observed in a long MD simulation of Cdc25B.^21,23^

**Figure 1:**
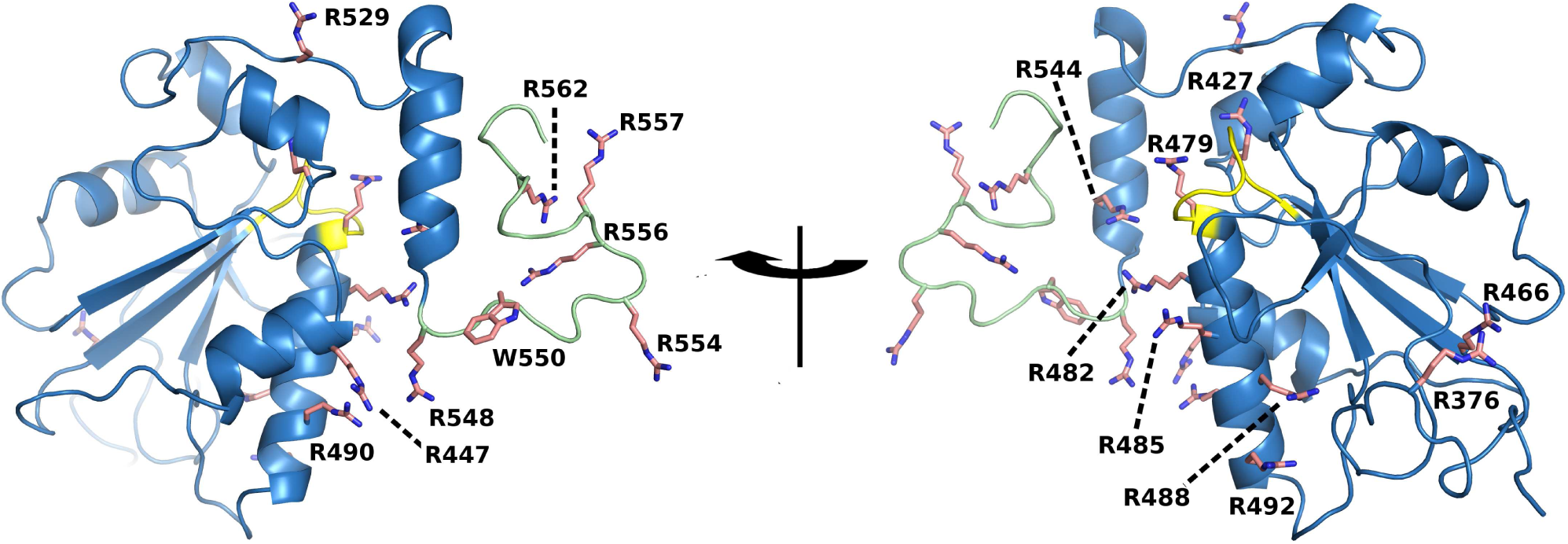
Structure of the complete catalytic domain of human phosphatase Cdc25B taken from a molecular dynamics snapshot and shown in cartoon with indicated Arg and Trp side-chains in pink sticks. The P-loop catalytic site is in yellow and the disordered C-terminal is in pale green.

In the following section the simulation and experimental methods are described with special care to assure the approximate force-field used in the protein simulations reproduces the cation-*π* energetics accurately. Then, free energy profiles estimated from umbrella sampling simulations are presented for the formation of eight putative cation-*π* contacts between side-chains of Trp-Arg pairs in Cdc25B. These results are validated by chemical shift perturbations obtained from NMR spectra of Cdc25B wild-type and point mutants disrupting putative cation-*π* contacts to identify important transient interactions in the protein conformational ensemble.

## 2 Materials and Methods

### 2.1 Quantum chemistry, molecular dynamics and free energy simulations

To evaluate the description of cation-*π* interactions by approximate force-fields, an isolated complex between blocked tryptophan (Ac-Trp-NHMe, denoted as bTrp) and blocked arginine (bArg) was used as a model system. The interaction energy with side-chains placed in two relevant orientations,^1^ sandwich and T-shape (Fig. 2), was evaluated quantum mechanically at the MP2/6-311++G** level corrected for the basis set superposition error with the counterpoise correction,^24^ and compared to CHARMM36^25^ and AMBER99-ILDN^26,27^ force-fields in the same geometries. The GAUSSIAN09 rev. A program was used for quantum calculations.^28^

**Figure 2:**
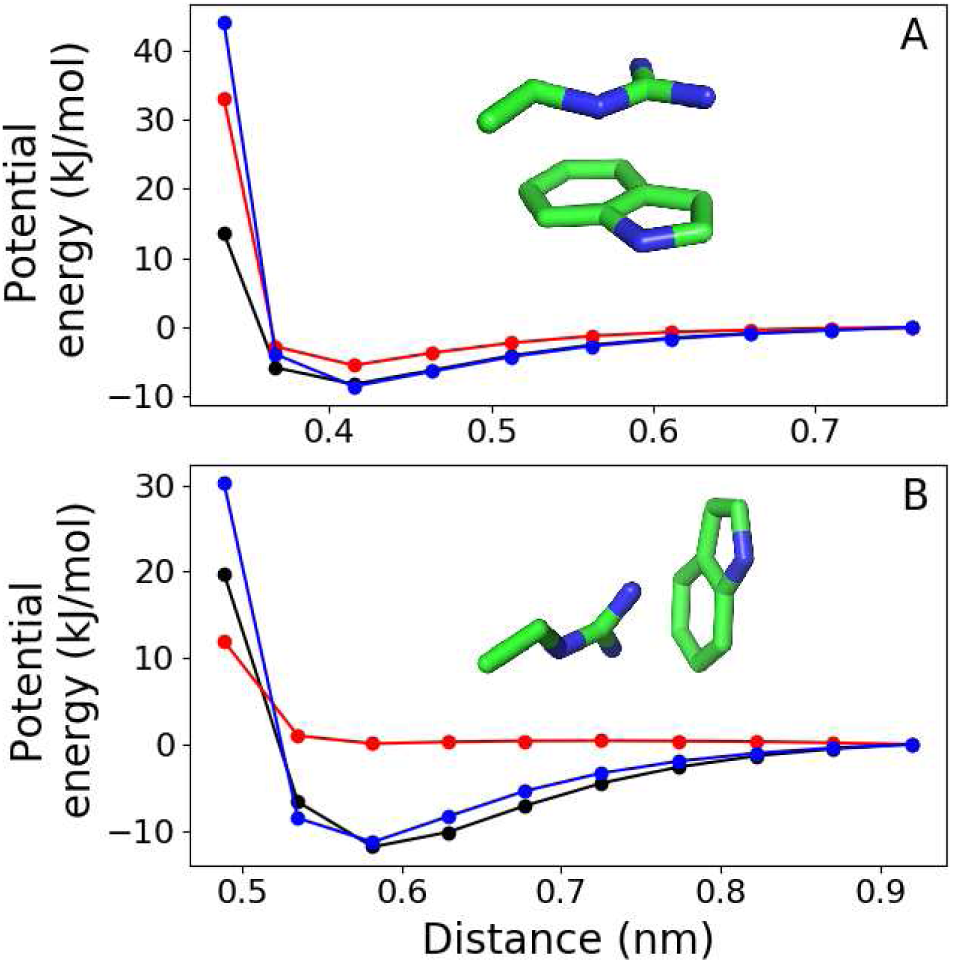
Potential energy for the interaction between Trp and Arg side-chains (bTrp · bArg complex) in sandwiched (panel A) and T-shaped (panel B) orientations, as shown in inserts. Black dots show the quantum chemical reference (MP2/6-311++G**), blue dots are the CHARMM36 force field and red dots show the AMBER99-ILDN force field. Lines are guides to the eye.

The reaction coordinate used here to describe cation-*π* interactions for both the bTrp · bArg complex and the Cdc25B free energy profiles below is the distance between the center-of-mass (COM) of all atoms in the Trp and in the Arg side-chains. These COMs roughly correspond to the position of Trp-C*δ*2 and Arg-N*ϵ* atoms.

Molecular dynamics simulations of the complete catalytic domain of the human phosphatase Cdc25B were initiated from the structure in PDB code 1QB0. Sixteen residues (A551-Q566) lacking from the native C-terminal structure^14^ and seven residues in the N-terminal part of the experimental construction (see below) were manually added in an extended conformation.^21^ Crystallographic water, ions and *β*-mercaptoethanol (BMER) were removed. Hydrogen atoms were added and a standard protonation state at pH=7 was assumed for side-chains. This is equivalent to the set-up we used previously.^21^ Two inorganic phosphate ions (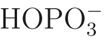form) were added manually to the catalytic P-loop (R479) and to the secondary phosphate binding site near R488 and R492 side-chains as previous data such as phosphate titrations strongly support that these sites will bind phosphate ions under our experimental conditions.^15,21,23^ This model was solvated in a dodecahedral simulation box of 13234 water molecules, with 4 Na^+^ and 2 Cl^−^ ions added to neutralize the system and mimic the experimental salt concentration. This system was relaxed and equilibrated during a 100 ns free MD simulation. The CHARMM36 force-field for protein and ions, and the TIP3P water model^29^ were used. Simulations were performed with a 2 fs integration step, 300 K temperature and 1 bar pressure (NPT ensemble) using the Bussi thermostat^30^ and Parrinello-Rahman barostat both with a period of 0.5 ps. PME method with 1.2 nm real space cutoff and 0.12 nm Fourier spacing was used to treat long-range electrostatics. All classical force field calculations, MD simulations and analysis were performed with the GROMACS 4.6.7 program.^31^

Free energy profiles were estimated with umbrella sampling (US).^32^ Initial configurations for each window were obtained after model equilibration described above. Seven US windows with reference reaction coordinate separated by 0.1 nm, from 0.4 to 1.0 nm, were used to estimate the profiles for each of Trp-Arg pair. A harmonic potential with force constant *k*_*umb,dCOM*_ = 2000 kJ mol^−1^ nm^−2^ was used. Each US window was sampled for 500 ns with a sample collected every ps. Thus, the total aggregate simulation time was 20 *µ*s. Potentials of mean-force were obtained with WHAM^32^ and the statistical uncertainty was estimated as 95% confidence intervals by bootstrap analysis with 50 resampling steps.^33^ The initial 50 ns of each window were discarded to allow equilibration of orthogonal degrees of freedom in all analysis presented. The overlap of reaction coordinate distribution between adjacent windows was confirmed.

### 2.2 Point mutants, protein expression and purification

Cdc25B W550A, R482A and R544A point mutants were generated from pET-28a plasmids with the complete wild-type Cdc25B catalytic domain (S373–Q566) cloned and fused to His-tag and thrombin cleavage sites. The N-terminal sequence after cleavage was GSHMEFQ (373)SDHRELI… where the first 7 residues are not native. Point mutations were introduced by using the QuickChange Lightning Site-Directed Mutagenesis kit (Agilent Tech. #210518).

Expression of wild-type Cdc25B and point mutants was made after *Escherichia coli* BL21-Gold (DE3) transformation with the respective plasmids and growth in M9-minimal (^15^NH_4_Cl-labeled) medium at 37°C until OD_600*nm*_ 0.6 (exponential growth phase). At this time, 0.8 mM isopropyl beta-thiogalactopyranoside (IPTG) solution was added for induction of protein expression, and temperature was maintained at 20°C for 18 hours when bacteria were then collected by centrifugation (7,000×*g*-20 min). Bacteria were lysed by sonication in lysis buffer (20 mM Tris-HCl, 500 mM NaCl, 10% Glycerol, 5 mM 2-mercaptoethanol (BMER), 1 mM phenylmethylsulfonylfluoride-PMSF, pH=7.4). The lysate was subjected to centrifugation and supernatant was subjected to column affinity (Ni-NTA) and size-exclusion (Superdex 75 HR 10/30) chromatographies to protein purification as previously described in detail.^21^

## 2.3 NMR spectra

All samples were prepared using 0.2 mM concentration of uniformly ^15^N-labeled proteins in the same phosphate buffer preparation (20 mM NaH_2_PO_4_, 50 mM NaCl, 5 mM BMER, 2 mM dithiotreitol, 5% ^2^H_2_O, pH=6.7) to obtain equivalent environmental conditions. NMR spectra were acquired on Bruker Ascend III 800 Mhz NMR apparatus with 4-channel TCI cryoprobe and Avance III console. Initial processing of the spectra was done with NMRPipe program^34^ and assignment transfer for the main and side-chain resonances was performed with CCPNMR Analysis Assign 2.4 program.^35^

The chemical shift perturbation (CSP) analyzed for each assigned N-H system was defined according to:

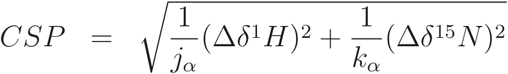

where Δ*δ* is the chemical shift change upon mutation, *j*_*α*_ and *k*_*α*_ are scaling factors (Table S1) that correspond to standard deviation of ^1^H and ^15^*N* chemical shifts of N-H system *α* (thus, unique to each amino-acid type, main and side-chains) as observed in the RefDB database.^36^ Statistical analysis was performed by the iterated removal of outliers defined as the values above the mean 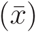 plus 3 standard deviations (*σ*).^37^ CSP were considered significant when larger than 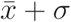.^38^

## 3 Results & Discussion

### 3.1 Benchmark of the empirical force field

The accuracy of a molecular simulation will depend on both energetic description and conformational sampling. Before addressing the importance of transient cation-*π* interactions in the protein Cdc25B, we bench-marked the interactions between Trp and Arg side-chains by comparing two widely employed protein force fields to high-level quantum chemical calculations.

Figure 2 shows the CHARMM36 force field correctly describes the cation-*π* interaction in both sandwiched (or stacking, parallel) and T-shaped (or perpendicular) orientations, with minimum energies within 0.5 kJ/mol of the MP2 energy reference. On the other hand, the AMBER99-ILDN energy function only agrees with the MP2 reference for the sandwiched orientation, with a difference of 3 kJ/mol for the minimum energy. Strikingly, the T-shaped orientation is fully non-binding with the AMBER force field. Thus, CHARMM36 was employed for US simulations presented below as this comparison suggests it should give a reliable energetic description for the mainly electrostatic interactions between Trp and Arg side-chains.^1^

Discrepancies between quantum and classical energy descriptions in small distances (0.3 nm for sandwiched and 0.5 nm for T-shaped orientations, Fig. 2) are due to the steep (and formally approximate) behavior of the repulsive component employed in force fields.^39^ This short-range discrepancy does not affect significantly the stability of cation-*π* interactions simulated here.

### 3.2 Free energy profiles for Trp-Arg cation-*π* interaction

Human Cdc25B catalytic domain contains 17 Arg, 13 Lys and only one Trp residue. We first analyzed the distance of each cationic residue to W550 in the Cdc25B x-ray structure^14^ and in a previous long MD simulation (total time of 6 *µ*s, Table S2)^21,23^ to check which cation-*π* interactions could be formed. We discarded Trp-Lys interactions and concentrated in Trp-Arg pairs, which interactions were shown above to be well described by the CHARMM36 force-field. Eight possible pairs were chosen and the free energy for interaction between their side-chains was estimated using umbrella sampling simulations. Results are shown in Fig. 3 and Fig. S1.

**Figure 3:**
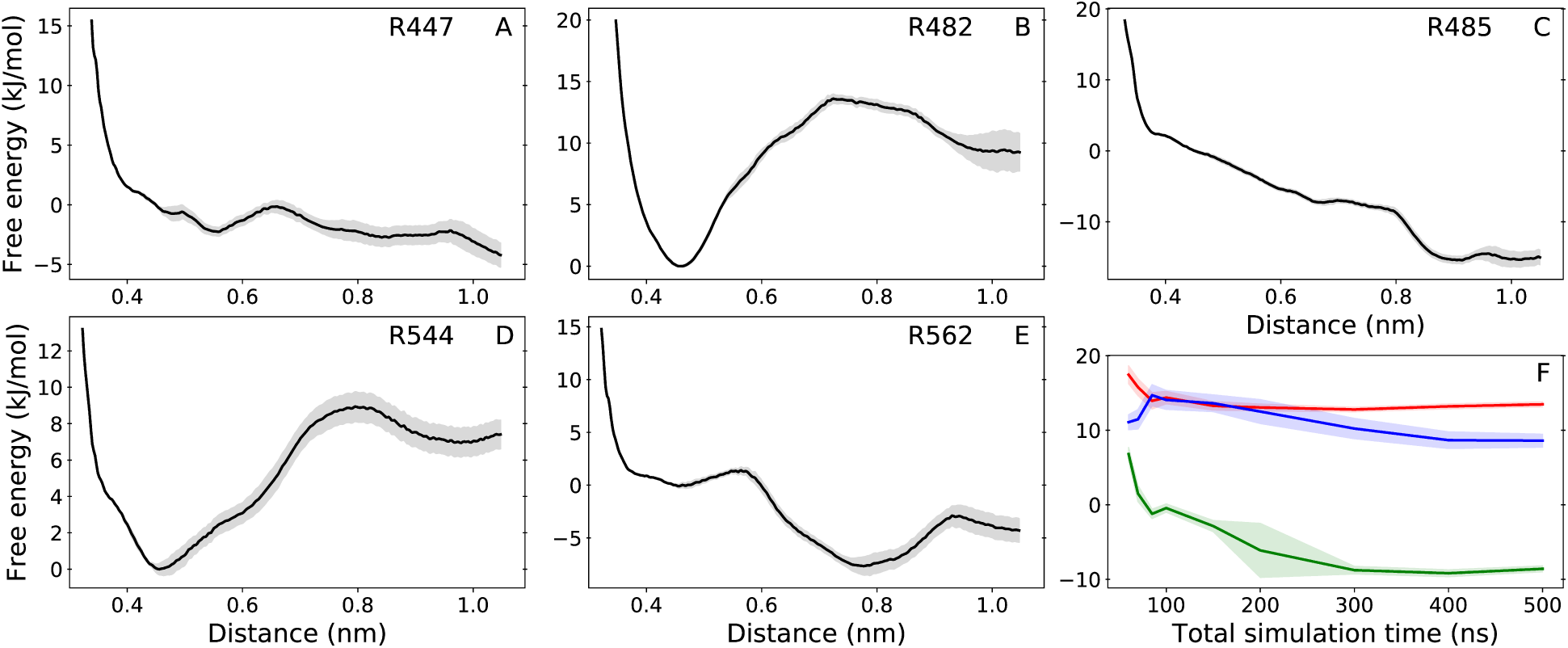
Free energy profiles for the cation-*π* interaction between W550 side-chain and R447 (panel A), R482 (B), R485 (C), R544 (D) and R562 (E) side-chains. Shadows around lines in all panels show the statistical uncertainty. Panel F shows convergence with total simulation time of the free energy difference between distance 0.45 and 0.80 nm for the cation-*π* complex with R482 in red line, R485 in green and R544 in blue.

There are three kinds of free energy profiles. For R482 and R544 (Fig. 3), the profile clearly shows a bound minimum at 0.45 nm distance and a dissociation barrier at 0.7-0.8 nm of 14 kJ/mol for R482 and 9 kJ/mol for R544, characterizing an important transient cation-*π* interaction with W550. For R485 (Fig. 3) and R490 (Fig. S1), the profile is downhill or dissociative, meaning that no stable interaction should be observed. For R447, R562 (Fig. 3), R554 and R556 (Fig. S1) the profile shows very weak minima and is almost flat in bound distances (0.4-0.8 nm), characterizing possible but short-lived and statistically unimportant interactions with W550.

The free energy difference between distance 0.45 and 0.80 nm in the profiles of Fig. 3, which corresponds to the dissociation barrier of bound profiles, converges to within one statistical uncertainty in less than 300 ns of total simulation time per US window (Fig. 3F). Thus, free energy profiles were estimated here with twice more sampling time than necessary for barrier convergence. Note the pair between W550-R548 was not studied in detail. Their neighbor side-chains will trivially engage in a cation-*π* contact, but it is not possible to validate this contact with the experimental analysis proposed below.

The minimum distance and dissociation barrier of bound profiles (Fig. 3B and D) are in good agreement with the distance and interaction energy of the minima for the bTrp · · · bArg complex found in our benchmark calculations (Fig. 2A). The distribution of the angle formed between the planes defined by the indole group of W550 and the guanidinum of R482 shown in Fig. 4A spreads out and shifts to higher angle values as the US reference co-ordinate assumes longer distances. The distribution is narrower in short distances because the T-shaped orientation is sterically hindered. In agreement with previous studies,^1^ a combination of mostly sandwiched but also T-shaped orientations is found for important transient cation-*π* interactions within the bound state (interval from 0.4 nm to 0.8 nm).

**Figure 4:**
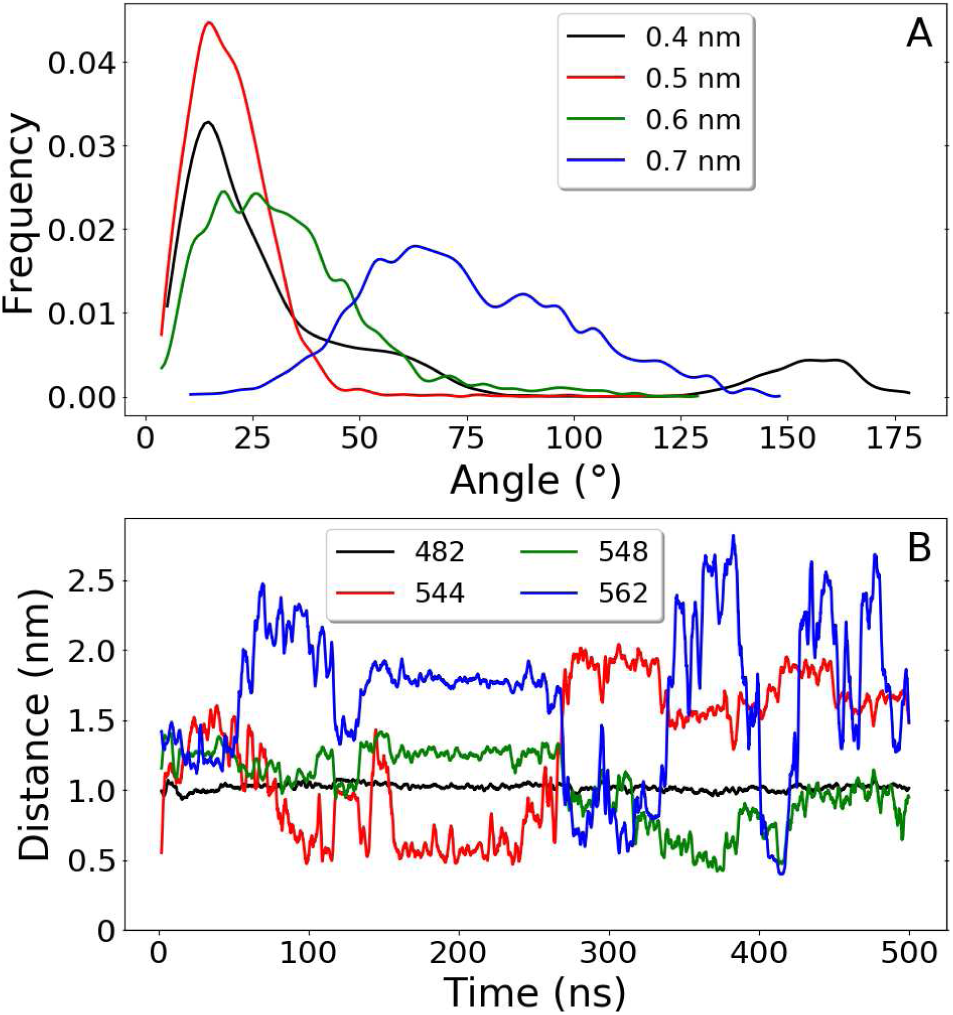
Analysis of Trp-Arg interactions during R482 US simulations. (A) Distribution of the angle formed between the planes defined by the indole group of W550 and the guanidinum of R482 with US reference coordinate centered in 0.4 (black), 0.5 (red), 0.6 (green) and 0.7 (blue) nm. Lower angles represent a sandwiched conformation and angles near 90° represent a T-shaped conformation, as in Fig. 2. Data were smoothed with a spline interpolation. (B) Distance trajectories between the COM of W550 side-chain and the COM of R482 (black), R544 (red), R548 (green) and R562 (blue) side-chains, obtained with US reference coordinate in 1.0 nm.

Given the large number of Arg residues in Cdc25B, many flanking W550 and the catalytic site, we asked which interactions would W550 perform when unbound from one of its important Arg partner. With R482 restricted to a long distance, Fig. 4B shows that W550 side-chain spontaneously forms complexes with R544 and R548, with very short-lived contacts with other Arg residues (R562 in this case). Similarly, when R544 is held away, R482 and R548 form complexes with W550 (data not shown). Thus, there is a competition of transient cation-*π* complexes With W550 in Cdc25B.

### 3.3 Validation by NMR chemical shift perturbation

We attempted to experimentally validate the importance of transient cation-*π* interactions by comparing standard ^1^H-^15^N HSQC spectra acquired for Cdc25B wild-type and point mutants W550A (Fig. S2), R482A (Fig. S3) and R544A (Fig. S4) in solution. Our premise was that a significant perturbation (*i.e.*, larger than 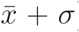) would be observed in the chemical shifts of residues performing important cation-*π* interactions by abolishing such contact upon a point mutation.

Main chain ^1^H-^15^N and Trp ^15^N*ϵ*_1_ side-chain signals were analyzed here as they are generally available and easy to acquire. We avoided additional NMR experiments (for instance, 2D HE(NE)HGHH)^12^ that would be necessary to assign Arg and Lys side-chain resonances. Possible environmental perturbations were minimized by acquiring all spectra in the same batch and buffer solution. Assignments were transferred for 144 peaks out of the 150 originally assigned^21^ and CSP was calculated for each point mutant, as shown in Table 1.

**Table 1:**
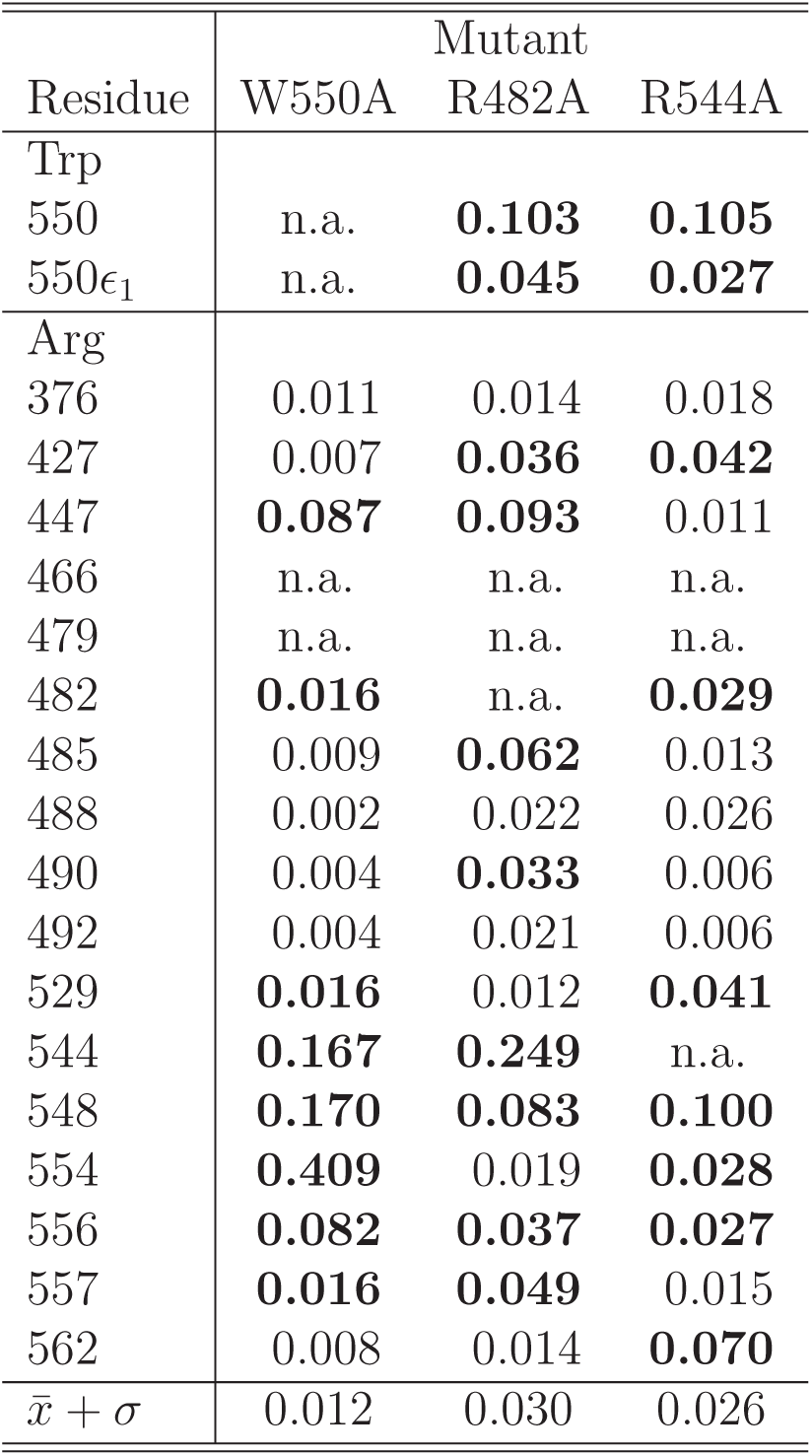
Chemical shift perturbation (CSP) of Arg and Trp main chain N-H centers for W550A, R482A and R544A point mutants of the Cdc25B phosphatase. The N*ϵ*_1_-H system in W550 side-chain is also shown. Unassigned centers are marked as n.a. Mean plus standard deviation (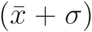) were calculated from around 115 CSP values for each point mutant after removal of outliers. Significative CSP are shown in bold.

| Residue | Mutant |  |  |
| --- | --- | --- | --- |
|  | W550A | R482A | R544A |
| Trp |  |  |  |
| 550 | n.a. | **0.103** | **0.105** |
| 550$\epsilon_1$ | n.a. | **0.045** | **0.027** |
| Arg |  |  |  |
| 376 | 0.011 | 0.014 | 0.018 |
| 427 | 0.007 | **0.036** | **0.042** |
| 447 | **0.087** | **0.093** | 0.011 |
| 466 | n.a. | n.a. | n.a. |
| 479 | n.a. | n.a. | n.a. |
| 482 | **0.016** | n.a. | **0.029** |
| 485 | 0.009 | **0.062** | 0.013 |
| 488 | 0.002 | 0.022 | 0.026 |
| 490 | 0.004 | **0.033** | 0.006 |
| 492 | 0.004 | 0.021 | 0.006 |
| 529 | **0.016** | 0.012 | **0.041** |
| 544 | **0.167** | **0.249** | n.a. |
| 548 | **0.170** | **0.083** | **0.100** |
| 554 | **0.409** | 0.019 | **0.028** |
| 556 | **0.082** | **0.037** | **0.027** |
| 557 | **0.016** | **0.049** | 0.015 |
| 562 | 0.008 | 0.014 | **0.070** |
| $\bar{x} + \sigma$ | 0.012 | 0.030 | 0.026 |

The CSP of cationic residues upon Trp mutation should be examined first. Insignificant perturbations found for R376, R427, R485, R488, R490, R492 and R562 in Table 1 imply that these residues do not perform important or long-lived cation-*π* interactions with W550. A similar conclusion is valid for most of the Lys residues (Table S3).

These observations are consistent with the Cdc25B crystal structure, where R376 and R427 are very distant from W550. But this flexible Trp residue approaches the other Arg located on the protein surface during a long MD simulation (Table S2). Why are their CSP insignificant? Side-chains of R488 and R492 make strong electrostatic interactions and remain buried within the secondary phosphate binding pocket.^15,21^ Side-chains of R485, R490 and R562 are fully exposed to solvent but do not perform a stable cation-*π* interaction with W550, as shown by the dissociative free energy profiles estimated here (Figs. 3 and S1).

A significant CSP is observed for R482 and R544 in the W550A mutant, in line with their bound free energy profiles. However, a significant CSP is also observed for R447, R529, R554, R556 and R557, in apparent disagreement with their almost flat free energy profiles and with the long distances to R529 and R557 (Table S2) that suggest no important interactions with W550 are formed. How can this be explained?

Chemical shifts are exquisitely sensitive to the distribution of protein conformations and to their interconversion rates.^11^ In Cdc25B, R554, R556 and R557 are close to W550 in sequence and placed in the same disordered C-terminal segment.^21^ If the W550A mutation alters the distribution or dynamics of main-chain configurations, it will lead indirectly to significant CSP for these Arg even if they do not perform important cation-*π* contacts. Although R447 and R529 are placed in the folded Cdc25B protein core, their segments have considerable flexibility as indicated by NMR order parameters (*S*^2^ ≤ 0.95 for D448-L453 and *S*^2^ ≤ 0.90 for D527-M531) and exchange rates (*R*_*ex*_ = 17 s^−1^ for E534) previously measured.^21^ Allosteric modulation of the flexibility along these segments by the W550A mutation may also lead indirectly to significant CSP.^40,41^

These indirect effects may also account for the significant CSP observed in protein regions close to mutation sites of R482A and R544A constructions, as well as for the CSP in R544 for the W550A mutant, although this Arg residue is in a stable *α*-helix. Thus, a significant CSP upon mutation of the *π*-system containing side-chain, here W550, is necessary but not sufficient to prove the importancy of a transient cation-*π* contact.

Mutations of cationic residues may also be explored. The CSP for W550 main-chain and ^15^N*ϵ*_1_ in the indole side-chain were significant for both R482A and R544A mutants (Table 1). This may be a relatively easy and general probe of transient cation-Trp interactions in other proteins as the Trp ^15^N*ϵ*_1_ signal usually appears isolated and can be readily identified even for unassigned HQSC spectra.^11^

Given that a competition of transient cation-*π* interactions is present (Fig. 4), abolishing one contact by mutation of the involved cationic residue would also directly perturb the molecular interactions (and the resonance signals) of the remaining Arg performing important contacts with the same *π* system. An indication of this competition in Cdc25B is the very high and localized CSP found for R544 upon R482A mutation (Fig. S3), even though these residues are structurally distant (Fig. 1). CSP is significant for R544 and R548 in the R482A mutant and for R482 and R548 in the R544A mutant. On the other hand, insignificant perturbations are found for R529 and R554 in the R482A mutant, and for R447 and R557 in the R544A mutant (R556 is off by 0.001 CSP units), indicating these five Arg do not perform important interactions with W550.

We may conclude that all CSP data (Table 1) and comparisons described here consistently suggest that *only* R482, R544 and R548 perform important transient cation-*π* interactions with W550 out of 15 assigned Arg residues in Cdc25B.

Despite the various structural and dynamical sources of CSP upon point mutations which complicate the identification of transient contacts mentioned above, another possible shortcoming of our NMR analysis is the non-unique formula used for calculation of CSP.^38^ Here we found essential to iteratively remove outliers^37^ and to scale chemical shift changes by their natural standard deviation obtained from a database.^36^ This is particularly relevant to be able to compare main-chain to side-chain CSP, as the dispersion of chemical shift for side-chain Trp-^15^N*ϵ*_1_ is much narrower than the dispersion of all main-chain shifts (Table S1).

## 4 Conclusions

Free energy profiles estimated from umbrella sampling with molecular dynamics simulations and perturbation of chemical shifts obtained from standard HQSC NMR experiments in solution were combined here to identify transient cation-*π* interactions of solvent-exposed Arg-Trp pairs in proteins. The phosphatase Cdc25B, containing several Arg residues and only one Trp located in a disordered segment, was used as a model system. Three kinds of free energy profiles were found: bound profiles (R482 and R544) with a clear dissociation barrier of 9-14 kJ/mol, characterizing an important transient cation-*π* interaction; dissociative profiles (R485 and R490), meaning that no cation-*π* interaction should be observed; and almost flat profiles (R447, R554, R556 and R562) with weak minima (*<* 5kJ/mol) in bound distances, characterizing possible but very short-lived and statistically unimportant cation-*π* interactions. The same three kinds of profiles should be found generally for transient cation-*π* interactions between solvent-exposed side-chains in other proteins.

All the free energy profiles obtained for eight putative cation-*π* pairs in Cdc25B are in qualitative agreement with the CSP analysis, confirming that R482 and R544 perform important transient interactions with W550 in solution, even though their side-chains are separated by more than 8 Å in the crystal structure. Although transient contacts of W550 with R447 and R485 were observed in previous MD simulations,^21^ results presented here clearly show that these contacts are not important in the Cdc25B conformational ensemble. Thus, the statistical significance of transient contacts in solution should not be relied upon single observations over crystal structures or molecular simulations.

The experimental validation proposed here could be generally applied to other proteins. It is relatively simple to conduct once the protein constructs are purified and the (wild-type) ^1^H-^15^N HSQC spectra is assigned. However, NMR chemical shits are sensitive to changes in both molecular contacts and conformational distributions. Disruption of a cation-*π* interaction upon mutation will result directly in a significant CSP by abolishing this molecular contact. But a significant CSP may also appear due to indirect changes in the residue conformational distribution. Thus, the proposed CSP analysis should be applied carefully and in combination with other statistically representative data.

## Acknowledgement

Funding from FAPESP (grants 2012/00543-1, 2016/24096-5, 2016/22365-9, 2018/25952-8 and 2018/08311-9), CNPq (fellowship 141683/2016-3 to A.A.O.R.), CAPES (Finance code 001) and computational resources from the SDumont cluster in the National Laboratory for Scientific Computing (LNCC/MCTI) are gratefully acknowledged.

## Supporting Information

**Table S1:** Standard deviation of ^1^H and ^15^N chemical shift (in ppm) for main chain N-H system of amino-acid type *α* obtained from the RefDB database.^36^

| Residue | Scaling factors |  |
| --- | --- | --- |
| | $j_\alpha$ ($^1\text{H}$) | $k_\alpha$ ($^{15}\text{N}$) |
| Ala | 0.660 | 4.040 |
| Arg | 0.640 | 3.960 |
| Asp | 0.710 | 4.460 |
| Asn | 0.600 | 4.070 |
| Cys | 0.730 | 4.331 |
| Gly | 0.760 | 3.730 |
| Glu | 0.640 | 3.670 |
| Gln | 0.620 | 3.730 |
| His | 0.730 | 4.090 |
| Iso | 0.750 | 4.460 |
| Leu | 0.700 | 4.131 |
| Lys | 0.680 | 3.851 |
| Met | 0.650 | 3.660 |
| Phe | 0.770 | 4.230 |
| Ser | 0.650 | 3.810 |
| Tyr | 0.790 | 4.550 |
| Thr | 0.650 | 4.941 |
| Val | 0.720 | 4.850 |
| Trp | 0.860 | 4.649 |
| Trp Nε1-H | 0.491 | 1.884 |

Note the ratio between $^1\text{H}$ and $^{15}\text{N}$ scaling factors ( $j_\alpha/k_\alpha$ ) is similar to 0.15, the standard value used in CSP studies,<sup>21,38</sup> for all N-H systems except for Trp N $\epsilon$ 1-H.

**Table S2:** Side-chain distance (in nm) to W550 in the crystallographic x-ray structure^14^ and minimum distance (in nm) observed along 6 *µ*s of MD simulation^21^ for all Arg residues of the complete Cdc25B C-terminal domain.

| Arg | X-ray | Long MD |
| --- | --- | --- |
| 376 | 3.71 | 1.85 |
| 427 | 2.07 | 1.27 |
| 447 | 1.25 | 0.37 |
| 466 | 3.81 | 2.26 |
| 479 | 1.47 | 0.54 |
| 482 | 1.19 | 0.29 |
| 485 | 2.10 | 0.33 |
| 488 | 2.62 | 0.66 |
| 490 | 2.02 | 0.31 |
| 492 | 2.84 | 0.46 |
| 529 | 2.32 | 2.07 |
| 544 | 0.83 | 0.52 |
| 548 | 0.97 | 0.30 |
| 554 | – | 0.30 |
| 556 | – | 0.40 |
| 557 | – | 0.80 |
| 562 | – | 0.36 |

**Table S3:**
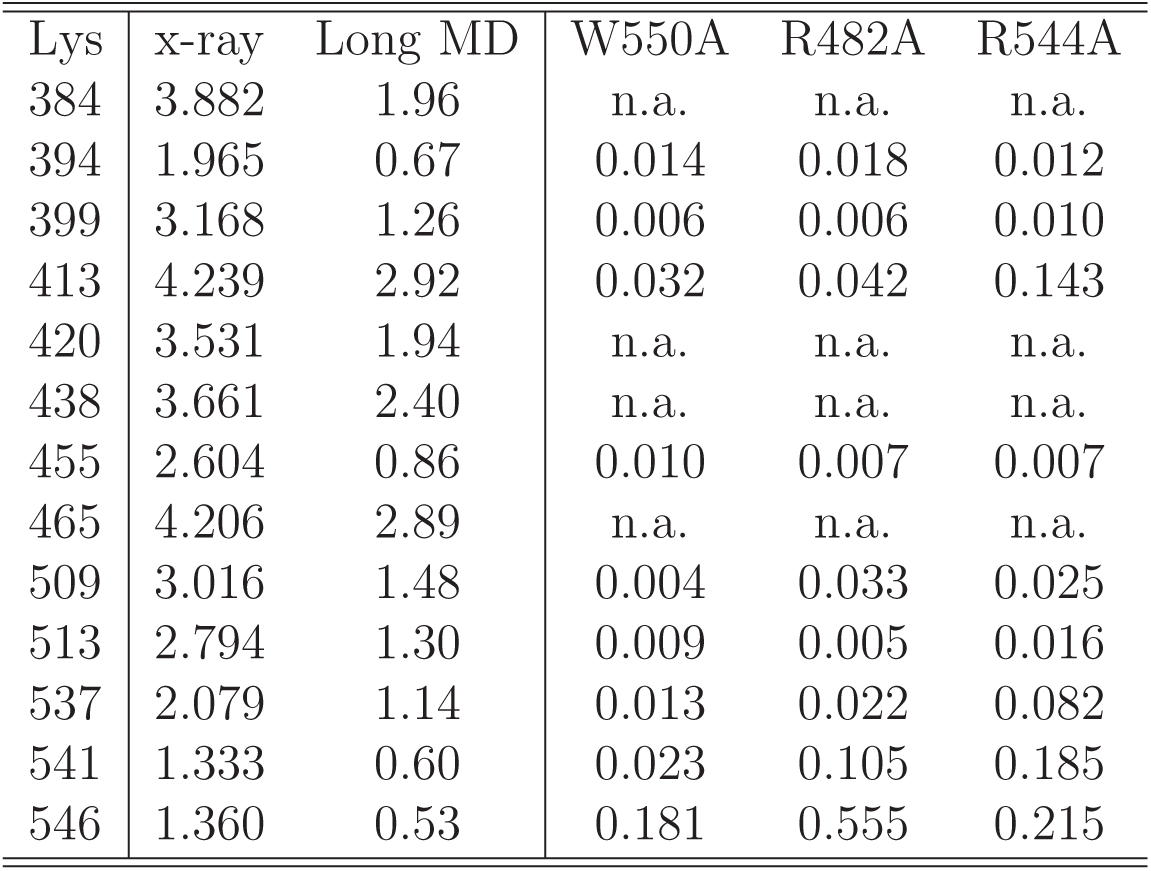
Side-chain distance (in nm) to W550 in the crystallographic x-ray structure,^14^ minimum distance (in nm) observed along 6 *µ*s of MD simulation,^21^ and chemical shift perturbation for W550, R482 and R544 point mutants of Cdc25B phosphatase, for all Lys residues of the complete C-terminal domain. Unassigned is marked n.a.

**Figure S1:**
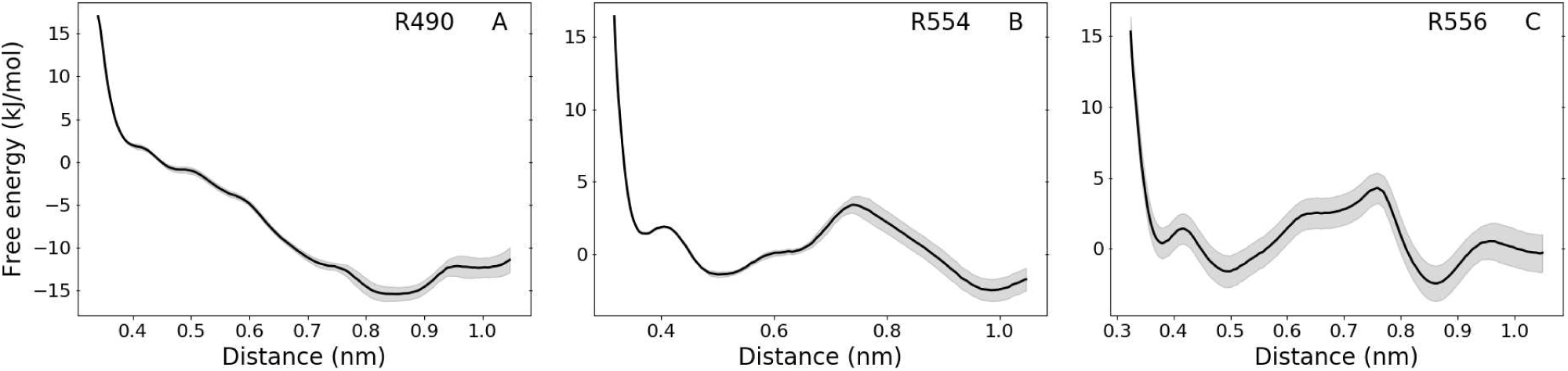
Free energy profiles for the cation-*π* interaction between W550 side-chain and Arg 490 (panel A), 554 (B) and 556 (C) side-chains. Shadows around lines in all panels show the statistical uncertainty. Profiles were obtained after 100 ns of total simulation time per US window and the first 30ns were discarded from analysis. These simulations were not carried out longer because the dissociative profiles shown and the low conservation of these Arg residues^21^ suggested these are not important interactions for the Cdc25B conformational ensemble.

**Figure S2:**
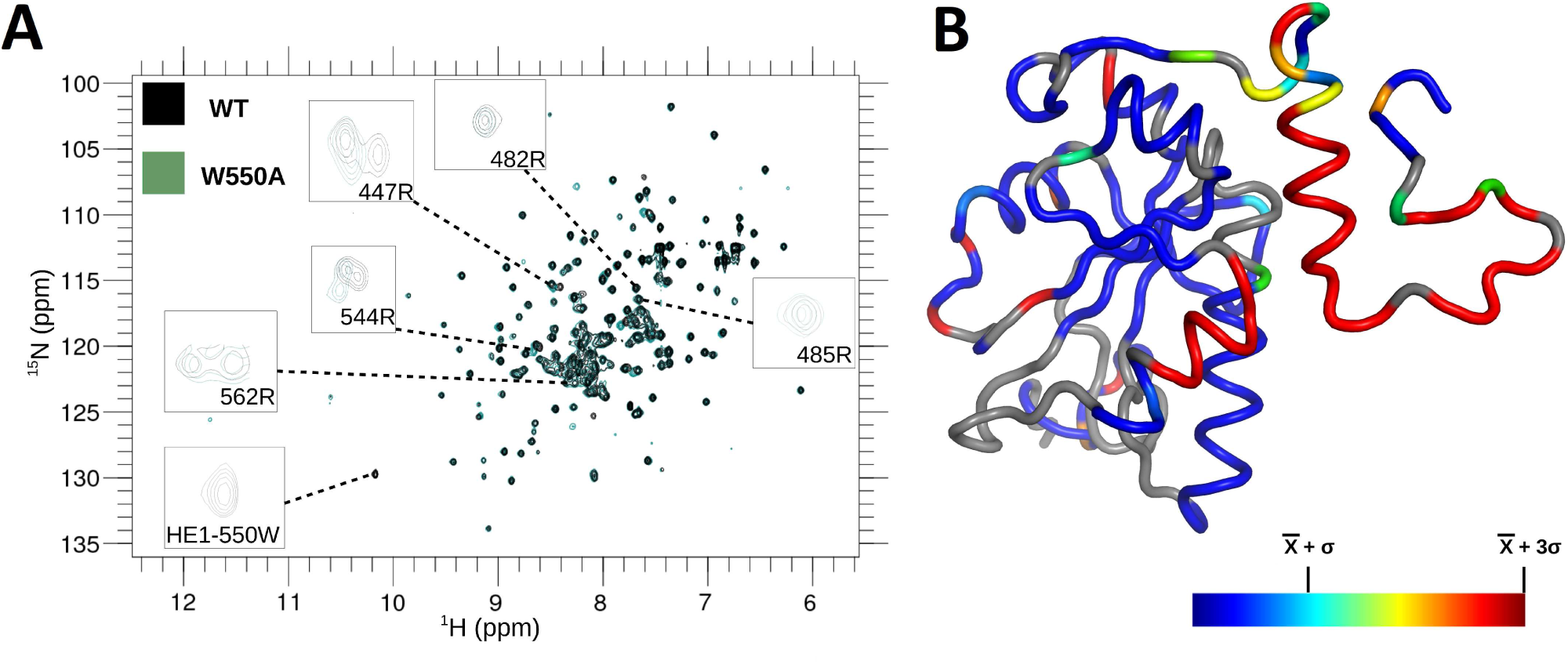
NMR spectra. (A) ^1^H-^15^N HSQC for Cdc25B wild-type and W550A point mutant and (B) chemical shift perturbation upon mutation projected according to colorbar over the same structure shown in Fig. 1. Gray segments were not assigned.

**Figure S3:**
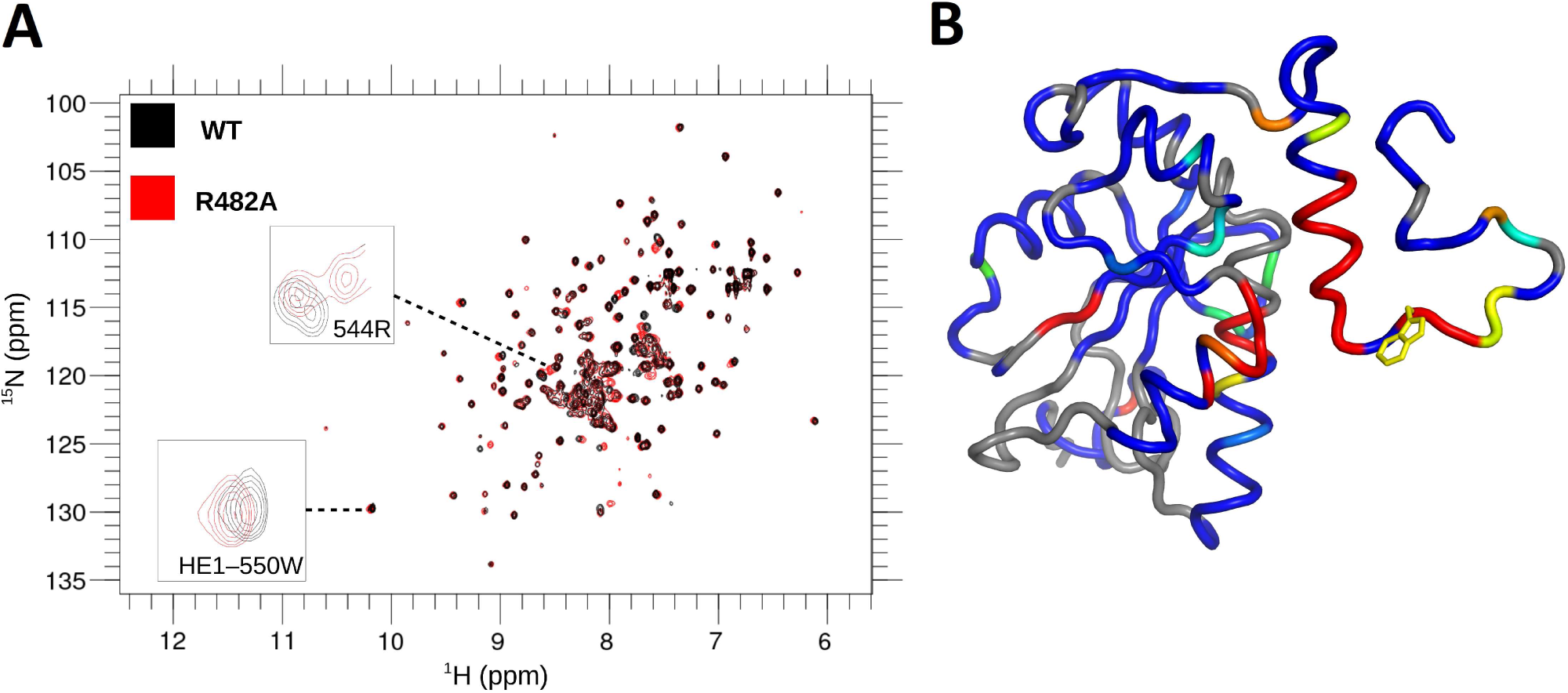
HSQC spectra and CSP as in Fig. S2 but for the R482A Cdc25B point mutant.

**Figure S4:**
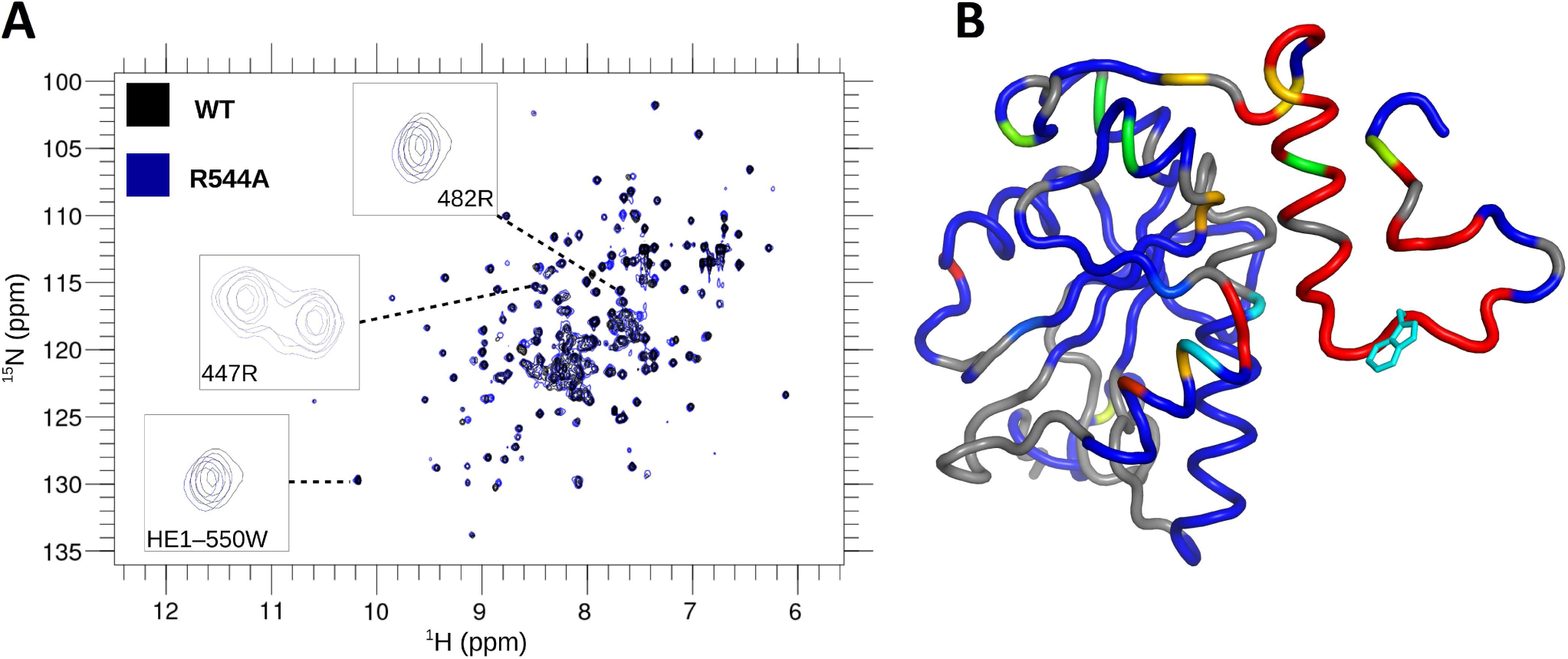
HSQC spectra and CSP as in Fig. S2 but for the R544A Cdc25B point mutant.

